# Quantitation, base identity and distribution of stably incorporated ribonucleotides in nuclear and mitochondrial DNA from murine tissues

**DOI:** 10.1101/2022.03.20.485043

**Authors:** Katrin Kreisel, Josephine Kalm, Sashidar Bandaru, Chandu Ala, Levent M. Akyürek, Anders R. Clausen

## Abstract

Ribonucleotides are estimated to be the most common non-canonical nucleotides transiently incorporated in DNA. Their presence or failure of their removal can affect genome stability and mutations in factors involved in dNTP pool maintenance or ribonucleotide removal can cause Aicardi-Goutières syndrome or promote certain human cancers. Here, we have mapped and quantitated ribonucleotides genome-wide, in nine tissues of wild-type mice. We observed tissue-specific variation in number and base identity of incorporated ribonucleotides and present evidence that a number of genomic features, such as tRNA genes, transcription start sites and G-quadruplexes, can increase the frequency of stably incorporated ribonucleotides in their proximity. Moreover, we present the non-random distribution of incorporated ribonucleotides in mtDNA and identified ribonucleotide hotspots. The study presents a framework to understand the physiological role of ribonucleotides in mammalian DNA.

## INTRODUCTION

After its discovery in 1869, DNA has been of great interest in research and was since then found to be the central hereditary molecule^1^. Without any known exceptions the genetic information of all eukaryotic life is stored in the form of DNA^2^. Despite the fact that some genetic diversity is beneficial from an evolutionary point of view in terms of population persistence and individual fitness^3^, DNA damage and mutations can have profound effects on the health and aging of an organism. It is therefore of great importance for each cell to maintain the integrity of its genetic material. The intactness of the DNA is however constantly challenged by intra- and extracellular factors: DNA damaging agents and irradiation have been investigated early on^4^, but more recently, knowledge of intracellular mechanisms affecting genome integrity is emerging^5^. Aside from the elucidation of DNA repair mechanisms, as well as error-prone and error-free translesion synthesis, a fundamental realization was the asymmetry of deoxyribonucleotide (dNTP) and ribonucleotide (rNTP) concentrations in the cell and later on in mitochondria^6,7^. Already in 1994 Thomas W. Traut compiled a list of nucleotide (NTP) concentrations illustrating in general the great excess of rNTPs as compared to dNTP concentrations and the variation of these concentrations between different cell types, tissues and organisms^6^.

Eukaryotic DNA is replicated by the three major replicative DNA polymerases α, δ and ε which discriminate against rNTPs with a so-called “steric gate” residue near the polymerases’ active site^8^. This discrimination however is not perfect and a direct consequence of the great rNTP excess is the relatively frequent misincorporation of ribonucleotides by replicative DNA polymerases. These events render ribonucleotides to be possibly the most common non-canonical nucleotides incorporated into the DNA^9^. While ribonucleotide incorporation may temporarily prove to be beneficial by mitigating challenging replication scenarios such as dNTP shortage^10^ and seem to fulfill useful functions such as allowing discrimination of parent and nascent strands during mismatch repair^11^ or serve as imprints for facilitating mating type switch in one of two *S. pombe* daughter cells^12^, their increased inclination to hydrolyze due to their 2’-hydroxyl group^13^ can cause nicked DNA and negatively affect genome stability^14^ resulting in associated pathologies^15,16^.

Since the incorporation of short RNA primers during the synthesis of Okazaki fragments is necessary and misincorporation of ribonucleotides by DNA polymerases happens frequently, as summarized by Sassa et al.^17^ mammalian cells possess multiple mechanisms to remove single ribonucleotides or short stretches of ribonucleotides from the DNA. RNA primers are removed either via DNA polymerase δ performing strand-displacement synthesis, displacing the primer from the previous Okazaki fragment, thereby forming a 5’-flap that is subsequently cleavage by flap endonuclease 1 and the resulting nick can be ligated by DNA ligase I^18^, or via RNA:DNA hybrid removal facilitated by the RNases H^19^. The removal of single ribonucleotides is mostly achieved by a mechanism called Ribonucleotide Excision Repair (RER) initiated by RNase H2, which cleave 5’ of embedded ribonucleotides in double-stranded DNA. Recent *in vitro* findings suggest a possible alternative RER pathway via the human DEAD-box RNA helicase DDX3X which showed RNase H2-like activity^20^. Topoisomerase 1 (TOP1) is also able to mediate the removal of a ribonucleotide but is more likely to generate short deletions at repetitive sequences or double-strand breaks^21,22^. It was hypothesized that incorporated ribonucleotides in yeast DNA may even be removed by Nucleotide Excision Repair as suggested by evidence in prokaryotes^23^, however findings in human model systems suggest that this DNA repair mechanism does not efficiently recognize incorporated ribonucleotides in the genomic DNA of higher eukaryotes^24^.

In contrast to RER and TOP1- or RNases H-mediated excision of ribonucleotides from nuclear DNA (nDNA), efficient ribonucleotide removal lacks in mitochondria. Mitochondrial DNA (mtDNA) underlies its own distinct replication and maintenance systems and mitochondrial NTP pools can affect mtDNA to a greater extend^25^, though increasing mtDNA instability with age has not been tied to its incorporated ribonucleotides^26^.

Genome-wide mapping of ribonucleotides has been used to track the usage of DNA polymerases^27^ and to determine how NTP pools affect the base identity and number of ribonucleotides incorporated into the nuclear and mitochondrial genome of human cell cultures and mouse liver^16,25,26^. Here, we hypothesize that there is variation in number and base identity of incorporated ribonucleotides when comparing different mouse tissues. We use the HydEn-seq method coupled with cleavage by the restriction enzyme SacI to map and quantitate stably incorporated ribonucleotides in the nuclear and mitochondrial DNA from nine different mouse tissues: blood, bone marrow, brain, heart, kidney, liver, lung, muscle and spleen. Since we use a wild-type mouse strain proficient in RER, this study for the first time gives a more comprehensive view on the ribonucleotide landscape within the DNA *in vivo*. We determined the frequency and base identity of incorporated ribonucleotides and provide evidence that in a wild-type mouse, ribonucleotide removal mechanisms are imperfect, and ribonucleotides remain permanently incorporated in both nDNA and mtDNA. Moreover, we provide evidence that incorporated ribonucleotides are not randomly distributed throughout the nuclear and mitochondrial DNA but are over- or underrepresented near certain genomic features in nDNA. Similarly, we observed non-random distribution of ribonucleotides in mtDNA with distinct hotspots throughout the molecule.

## RESULTS

Based on the knowledge that dNTP and rNTP pools vary between tissues^6^ and that their concentrations may affect how many and which ribonucleotides are misincorporated into the DNA^25^, we hypothesized that incorporated ribonucleotides vary in number and base identity in different mouse tissues. To test this hypothesis, we used the HydEn-seq method coupled with SacI digestion for the mapping and quantitation of incorporated ribonucleotides in nine tissues (blood, bone marrow, brain, heart, kidney, liver, lung, muscle and spleen) from six male wild-type mice. For quantitation of incorporated ribonucleotides, two HydEn-seq libraries were prepared for each sample: Both samples were cleaved with SacI, then one underwent alkaline hydrolysis with KOH and one was treated with KCl instead. All reads at 5’-ends were normalized to the mean reads in the SacI cleavage sites and reads at 5’-ends from KCl-treated libraries were subtracted from the corresponding KOH-treated libraries in order to determine the number of reads corresponding to incorporated ribonucleotides alone. Furthermore, the number of incorporated ribonucleotides was normalized to 1 kb for comparison.

### Number of embedded ribonucleotides varies between nDNA and mtDNA and is tissue-dependent

In nDNA (Fig. 1 A), we determined that approximately 1 ribonucleotide per kb was present and the variation in numbers of permanently incorporated ribonucleotides between tissues was relatively small with kidney and liver containing the least and spleen, lung and heart containing the most incorporated ribonucleotides. As hypothesized based on the lack of efficient ribonucleotide removal in mtDNA (Fig. 1 B), we observed more variance between tissues with approximately 1 to 8 stably incorporated ribonucleotides per 1 kb in mtDNA, with lung, spleen and muscle containing markedly more ribonucleotides than the other tissues. It is conceivable that tissue-specific differences in ribonucleotide incorporation exist even in nDNA, but that such differences are masked by the presence of efficient ribonucleotide removal mechanisms in the nucleus. To investigate the variation in the number of incorporated ribonucleotides among tissues, we performed the Welch’s t-test and reported the p-values in matrices. As evident from Figure 1 C and D, with several p-values below 0.05 the number of embedded ribonucleotides is statistically significantly different between multiple tissues: In nDNA (Fig. 1 C), the number or incorporated ribonucleotides did not vary greatly, but statistically significant differences were found between spleen and all tissues except lung and blood, and among kidney, liver and heart, which all showed little variation within the sample.

**Figure 1.**
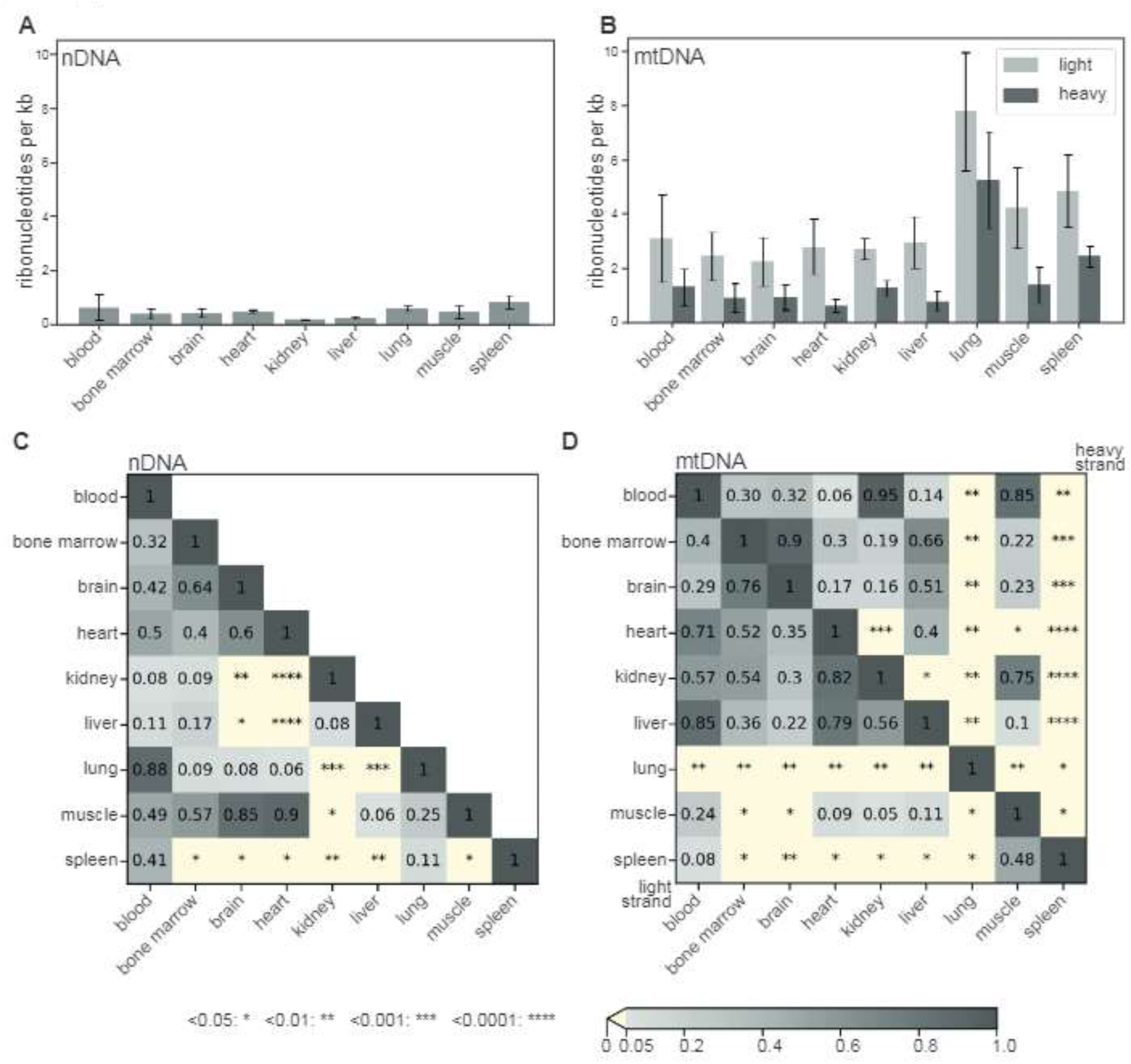
Genome wide quantitation of ribonucleotides in nDNA and mtDNA. **A** Mean ribonucleotides per kb in nDNA from blood, bone marrow, brain, heart, kidney, liver, lung, muscle and spleen. Error bars represent the standard deviation (SD). **B** Mean ribonucleotides per kb on heavy (dark grey) and light (light grey) strand of mtDNA from blood, bone marrow, brain, heart, kidney, liver, lung, muscle and spleen. Error bars represent the SD. **C** p-values of the difference between mean ribonucleotides per kb in nDNA from each tissue were calculated using the Welch’s t-test. **D** p-values of the difference between mean ribonucleotides per kb on the light (lower left matrix half) or the heavy strand (upper right matrix half) of mtDNA from each tissue were calculated using the Welch’s t-test. (n = 6 for blood, brain, heart, kidney, liver and muscle; n=5 for bone marrow, lung and spleen)

On both strands of mtDNA (Fig. 1 D) from lung and spleen the number of incorporated ribonucleotides was statistically significantly higher than in other tissues. The number of incorporated ribonucleotides varies not only between tissues (Fig. 1 A-D) but in case of mtDNA also between strands (Fig 1 B), presumably due to the unequal base composition (Fig. 2 C) between the two strands. We found the light strand to consistently contain about 1.5 to 2-fold more incorporated ribonucleotides than the heavy strand. Taken together, these results show that ribonucleotides are frequently present with around 1 ribonucleotide per kb in nDNA or approximately 5.2 million ribonucleotides per nuclear mouse genome, despite efficient methods of ribonucleotide removal and that ribonucleotides in mtDNA are more frequent than in nDNA with about 1 to 8 ribonucleotides per kb or 16 to 128 ribonucleotides per mtDNA molecule depending on the tissue, probably due to the lack of efficient removal mechanisms. The number of permanently incorporated ribonucleotides is moreover varying depending on the tissue in both nDNA and mtDNA.

**Figure 2.**
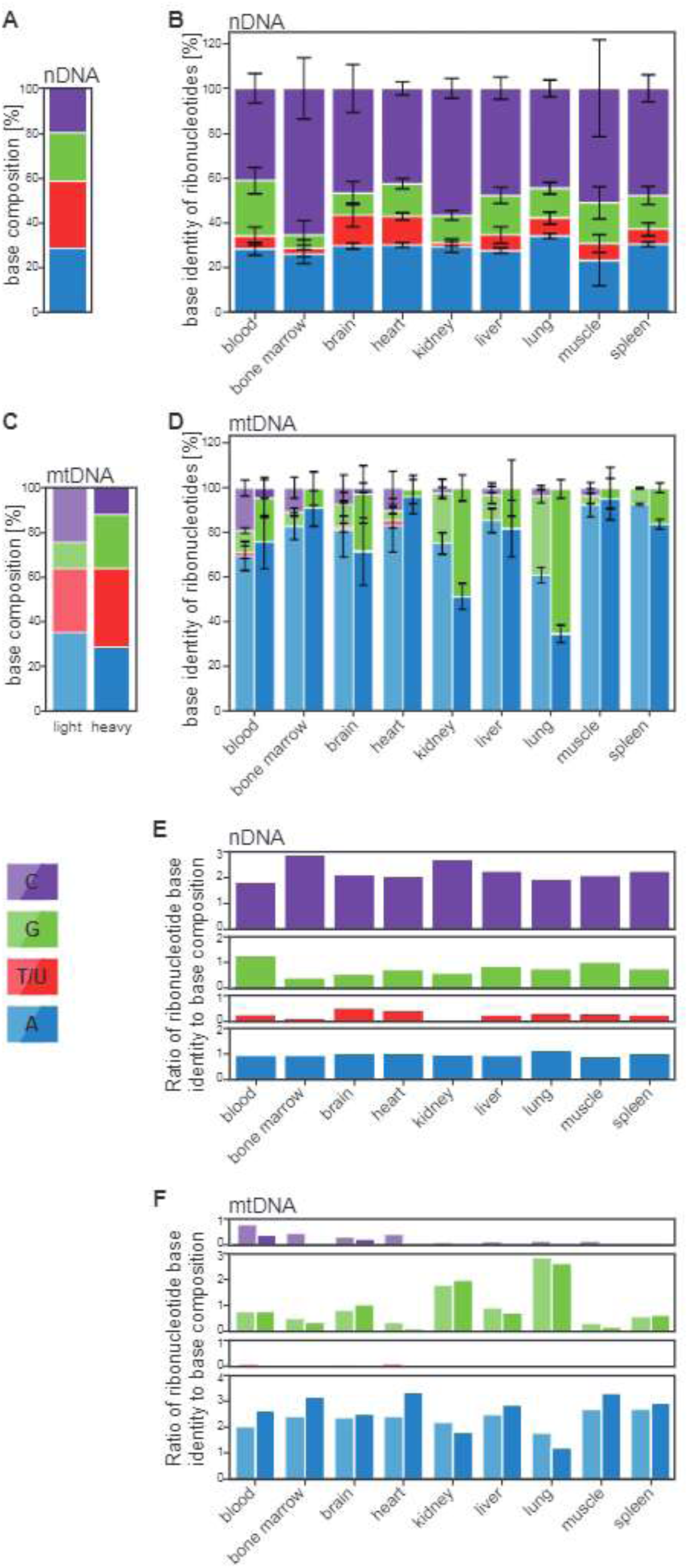
Base identity of incorporated ribonucleotides between tissues and nDNA and mtDNA. **A** Base composition of murine nDNA. **B** Ribonucleotide base identity in nDNA from blood, bone marrow, brain, heart, kidney, liver, lung, muscle and spleen. Error bars represent the standard deviation (SD). **C** Base composition of light and heavy strand of murine mtDNA. **D** Ribonucleotide base identity on the light (light shades) or heavy strand (dark shades) of mtDNA from blood, bone marrow, brain, heart, kidney, liver, lung, muscle and spleen. Error bars represent the SD. **E** Ribonucleotide enrichment was calculated by normalizing the ribonucleotide content with the nDNA base composition. **F** Ribonucleotide enrichment was calculated by normalizing the ribonucleotide content on light and heavy strand with the light or heavy strand base composition, respectively. (n = 6 for blood, brain, heart, kidney, liver and muscle; n=5 for bone marrow, lung and spleen)

### The base identity of incorporated ribonucleotides shows tissue-specific variation

Variation between tissues is not only reflected in the total numbers of incorporated ribonucleotides, but in the base identity of these ribonucleotides. Figure 2 A and C display the base composition of nDNA and mtDNA, respectively. As shown in Figure 2 B and D, the base identity of incorporated ribonucleotides differs from the base composition and the ratios between the four possible bases vary between tissues, as well. In nDNA, cytidine monophosphate (rC) followed by adenosine monophosphate (rA) were most commonly present in all samples, and to a lesser extend guanosine monophosphate (rG) and little to no uridine monophosphate (U) were observed. In contrast, rA was by far the most commonly incorporated ribonucleotide in mtDNA followed by varying amounts of rG, little to no rC and almost no U. Furthermore, we calculated the enrichment of incorporated ribonucleotides by normalizing the percentages of base identity to the base composition of nDNA (Fig. 2 E) and mtDNA (Fig. 2 F). If the incorporated ribonucleotides were proportionate to the base composition, values of 1 would be expected. Figure 2 E illustrates that rC is more frequently present than rG and rA with values around 1, while U is underrepresented in nDNA. In mtDNA (Fig. 2 F) we find rA to be preferentially incorporated, followed by rG which was overrepresented in kidney and lung, as well. In contrast, rC occurred rarely, and the presence of U was minimal in mtDNA. The high frequency of rA incorporation in mtDNA might also in part explain the higher total of incorporated ribonucleotides observed on the light strand which contains more adenine than the heavy strand (Fig. 2 C). The difference between nDNA and mtDNA samples may be the result of multiple differences between nucleus and mitochondria: Since nDNA and mtDNA are synthesized by different replicative DNA polymerases, DNA polymerases α, δ, ε in the nucleus and DNA polymerase γ in mitochondria, the architecture of each DNA polymerase affects which type of ribonucleotides are most likely incorporated by them and how frequently^9^. Furthermore, ribonucleotide repair mechanisms may have varying removal efficiency depending on the base identity and sequence context, possibly obscuring biases in incorporation of the nuclear DNA polymerases. As shown earlier in mtDNA, ribonucleotide incorporation is also directly affected by the rNTP and dNTP pools^25^. Consistent with earlier findings in mouse liver, heart and brain^16^ and rNTP and dNTP pool measurements in mouse liver mitochondria that showed an abundance of adenosine triphosphate (ATP)^28^, we found rA to be most commonly incorporated in mtDNA, with the highest percentage of incorporated rA in muscle mtDNA. Overall, these results suggest that the base identity of stably embedded ribonucleotides has tissue-specific variation in both nDNA and mtDNA and strengthens earlier evidence of the absence of ribonucleotide repair mechanisms^29^ and the high ATP concentrations in mitochondria to be the main reason for frequent rA presence in mtDNA.

### Ribonucleotide distribution in the genome shows distinct patterns at multiple genomic elements

We hypothesized that the distribution of ribonucleotides is not random and that the presence of genomic elements can affect the efficiency of ribonucleotide incorporation and removal for example by masking incorporated ribonucleotides from RER. To test our hypothesis that incorporated ribonucleotides remain more often unrepaired near genomic elements as compared to random positions in the genome, we performed meta-analyses where we investigated the number of ribonucleotides incorporated in bins located before and after genomic elements of interest. We obtained lists of genomic positions for enhancers^30^, tRNAs^31^, promoters^32^, CpG islands^33^ and microsatellites^34^ from the UCSC table browser^35^, transcription start sites (TSS) from refTSS^36^ and predicted intrastrand G-quadruplexes (G4s) by using a custom script searching for the pattern (G_3+_N_1-25_)_3_G_3+_ in the *Mus musculus* reference genome (mm10/GRCm38).

We filtered our ribonucleotide mapping data by the positions listed for each region of interest (ROI) and then calculated the mean number of incorporated ribonucleotides in bins before and after the ROI start positions. Enrichment plots (Fig. 3) were generated by dividing the mean number or incorporated ribonucleotides at ROI by the mean number from a random data set subjected to the same normalizations and binning. The random data set was generated by picking 1,000 random positions per chromosome. Display window and bin sizes were adjusted in relation to the ribonucleotide patterns observed at ROIs and spanned either 10 kb, 4 kb, 1 kb or 500 b up- and downstream of the ROI start position with 500 b-, 200 b-, 50 b-, or 25 b-bins, respectively. The analysis was performed separately for the same and opposite strand if strand information was available, otherwise reads on both strands were counted.

**Figure 3.**
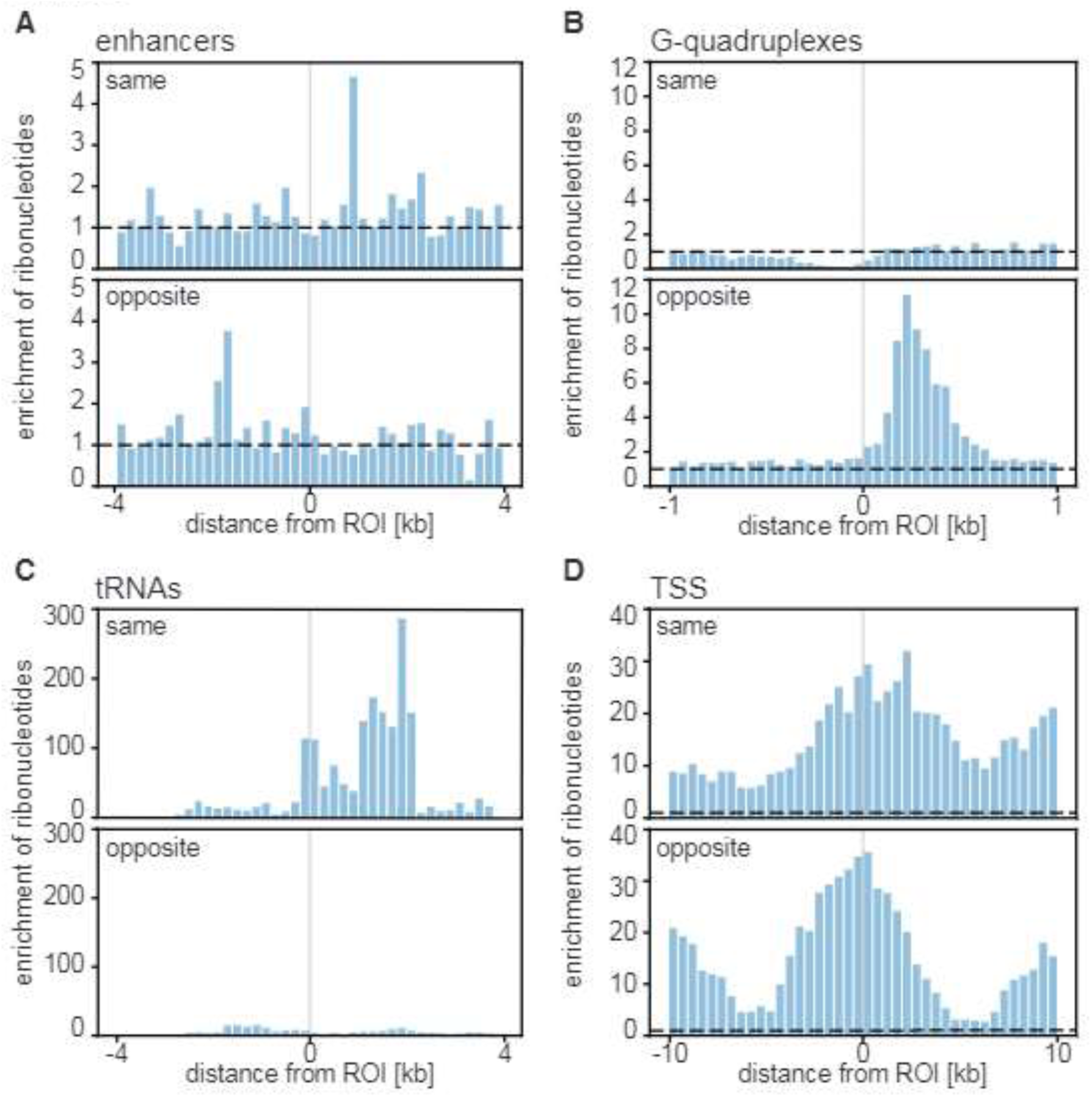
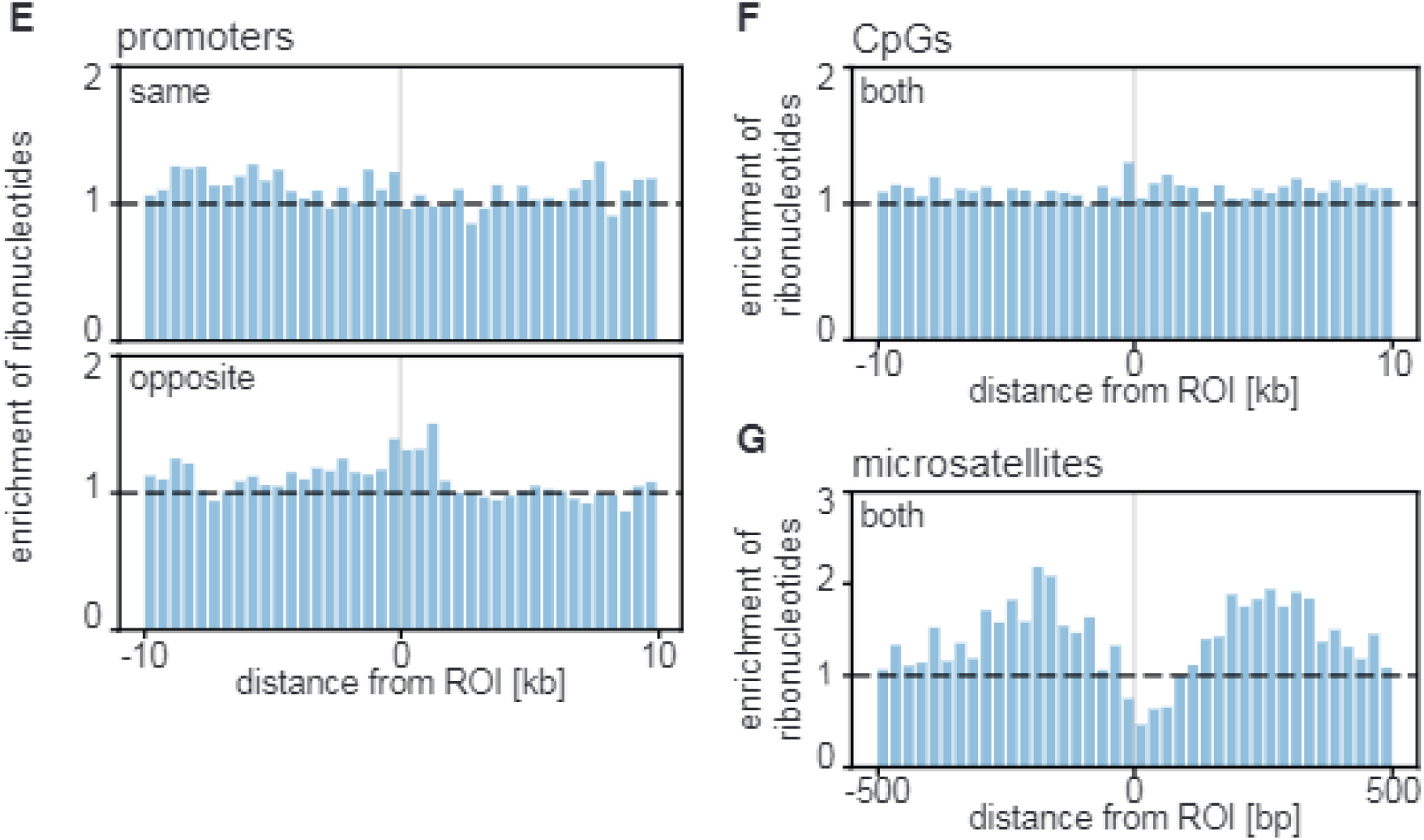
Enrichment of ribonucleotides at genomic features. Enrichment of ribonucleotides near genomic region of interest (ROI) on same, opposite or both strands. **A** Enrichment of ribonucleotides near 863 enhancers on the same and opposite strands. Mean ribonucleotide enrichment per 200 b-bin in a 4 kb-window up- and downstream of the ROI is shown. **B** Enrichment of ribonucleotides near 1,048,573 intrastrand G-quadruplexes on the same and opposite strands. Mean ribonucleotide enrichment per 50 b-bin in a 1 kb-window up- and downstream of the ROI is shown. **C** Enrichment of ribonucleotides near 435 tRNAs on the same and opposite strands. Mean ribonucleotide enrichment per 200 b-bin in a 4 kb-window up- and downstream of the ROI is shown. **D** Enrichment of ribonucleotides near 173,204 transcription start sites (TSS) on the same and opposite strands. Mean ribonucleotide enrichment per 500 b-bin in a 10 kb-window up- and downstream of the ROI is shown. **E** Enrichment of ribonucleotides near 25,111 promoters on the same and opposite strands. Mean ribonucleotide enrichment per 500 b-bin in a 10 kb-window up- and downstream of the ROI is shown. **F** Enrichment of ribonucleotides near 16,023 CpG islands on both strands. Mean ribonucleotide enrichment per 500 b-bin in a 10 kb-window up- and downstream of the ROI is shown. **G** Enrichment of ribonucleotides near 197,237 microsatellites on both strands. Mean ribonucleotide enrichment per 25 b-bin in a 500 bp-window up- and downstream of the ROI is shown. (All tissues, n = 51)

Our analysis of 863 enhancers (Fig. 3 A) showed a 4 to 5-fold enrichment of ribonucleotides at about 0.8 to 1 kb on the same strand as the enhancers and a nearly 4-fold enrichment at about -2 kb and - 1.6 kb on the opposite strand. Investigating 1,048,573 intrastrand G4s (Fig. 3 B), we found a depletion of ribonucleotides from about -500 b upstream to the start position of the G4 motif on the same strand while the opposite strand showed a steep, 10-fold enrichment of ribonucleotides in the bins from 0-500 b, peaking between 200 b and 300 b downstream of the start position of the intrastrand G4s. We analyzed 435 tRNAs (Fig. 3 C) and found that ribonucleotides were enriched by about 100-fold in a 200 b-window up- and downstream on the same strand as the tRNA starting position and further downstream peaking with a nearly 300-fold enrichment about 2 kb downstream, while we did not find any marked increase on the opposite strand. When analyzing 173,204 TSS (Fig. 3 D) and found a 30-to 35-fold enrichment of ribonucleotides near the TSS start point, decreasing to about 5-fold enrichment at around 5 kb up- and downstream, then increasing again in the periphery. Given that tRNA genes are short, we suspect that the peripheral increase in incorporated ribonucleotides may be a result of neighboring features that also show enrichment of ribonucleotides. We could not decern a clear pattern for the 25,111 promoters (Fig. 3 E), though there might be a slight overrepresentation of ribonucleotides on the opposite strand near the promoter start position. Since CpG islands and microsatellites are genomic features on both strands, we calculated the enrichment of ribonucleotides on both strands. Our analysis of 16,023 CpG islands (Fig. 3 F) did not show any noteworthy enrichment or depletion of ribonucleotides. The analysis of 197,237 microsatellites (Fig. 3 G) yielded an interesting trend, where at 500 bp up- and downstream of the microsatellite starting point the ribonucleotides are neither enriched nor depleted, then increase symmetrically by about 2-fold at 250 bp up- and downstream to then markedly decreased below the expected base level of incorporated ribonucleotides by 0.5-fold at the start position of the microsatellites. Taken together, these analyses suggest that ribonucleotides stay more frequently stably embedded at and around certain genomic features, warranting further investigation as to whether they are the result of increased incorporation or inhibited removal.

### Embedded ribonucleotides in mtDNA are located at distinct hotspots

We calculated the mean ribonucleotides at each position of the heavy and light strand of the mtDNA. As shown in Fig. 4 A for the heavy strand and Fig. 4 B for the light strand, the distribution of ribonucleotides is heterogenous and ribonucleotides are found at distinct hotspots throughout each strand. To best fit the data, the upper bound for the y-axis was set to 0.003 ribonucleotides per heavy strand and 0.005 ribonucleotides per light strand. Based on the quantitation in Fig. 1 B, estimating the average total number of ribonucleotides on the light strand to be around 3.5 which equates to about 0.0002 ribonucleotides at each position, and on the heavy strand around 1.6 ribonucleotides or circa 0.0001 ribonucleotides at each position, it is evident that certain positions are more prone to contain ribonucleotides as they show 10-to 20-fold higher numbers.

**Figure 4.**
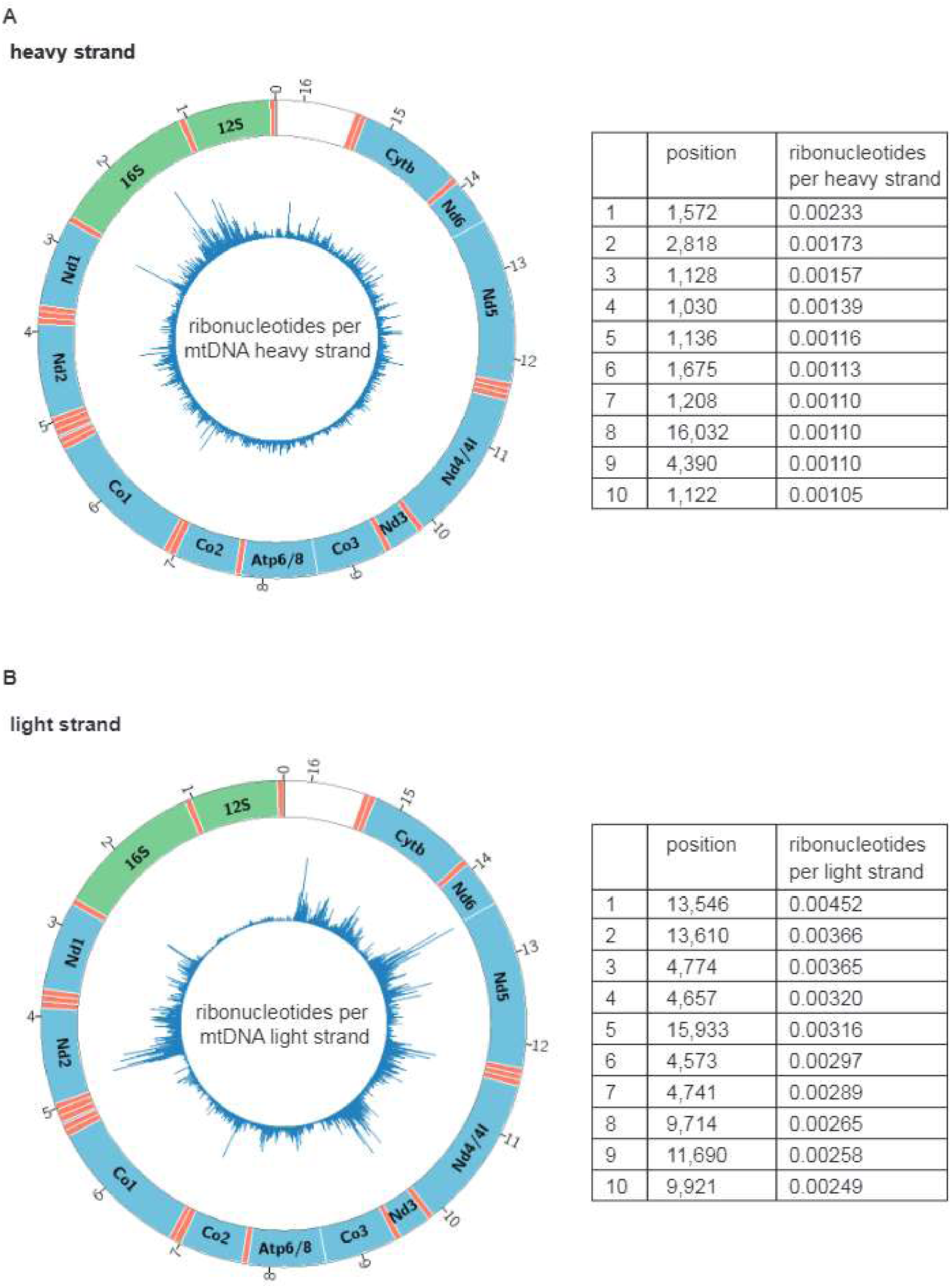
Distribution of ribonucleotides mtDNA. Genomic positions on the mtDNA are indicated in kb on the outer ring and coding regions (green, blue) were labeled with gene symbols. tRNAs are displayed as orange bands. **A** Circos plot (left panel) of incorporated ribonucleotides per heavy strand. The upper bound of the y-axis was set to 0.003 ribonucleotides per heavy strand to fully display the data. Top 10 ribonucleotide hotspots on the light stand (right panel). **B** Circos plot (left panel) of incorporated ribonucleotides per light strand. The upper bound of the y-axis was set to 0.005 ribonucleotides per light strand to fully display the data. Top 10 ribonucleotide hotspots on the heavy stand (right panel). (All tissues, n = 51)

Owing to the presence of RNase H1 in the mitochondria^37^, RNA primers synthesized by the mitochondrial RNA polymerase^38,39^ seem to be removed efficiently whereas single or diribonucleotides, as incorporated by DNA polymerase γ, seem to lack efficient removal pathways^25,29^. Based upon this, we hypothesized that ribonucleotides are underrepresented near the origin of heavy strand replication (OriH) and the origin of light strand replication (OriL). Surprisingly, we observed a distinct ribonucleotide peak at OriH at position 16,032 on the heavy strand (Fig. 5 A and B) with 0.001 ribonucleotides per heavy strand, corresponding to a 10-fold increase compared to the average if the distribution of the observed ribonucleotides was random. Likewise, we observed a peak at OriL (Fig. 5 C and D) at position 5,188 on the light strand with 0.001 ribonucleotides per light strand, corresponding 20-fold increase compared to the expected average number of ribonucleotides per light strand, if the distribution was random. In both cases the vicinity did not show a note-worthy enrichment in ribonucleotides. It is conceivable that the observed ribonucleotides at 16,032 and 5,188 are the result of imperfect primer removal, whether this is indeed the case or whether they were misincorporated later on which in general seems to be the main pathway for ribonucleotide incorporation^40^, however, remains to be determined.

**Figure 5.**
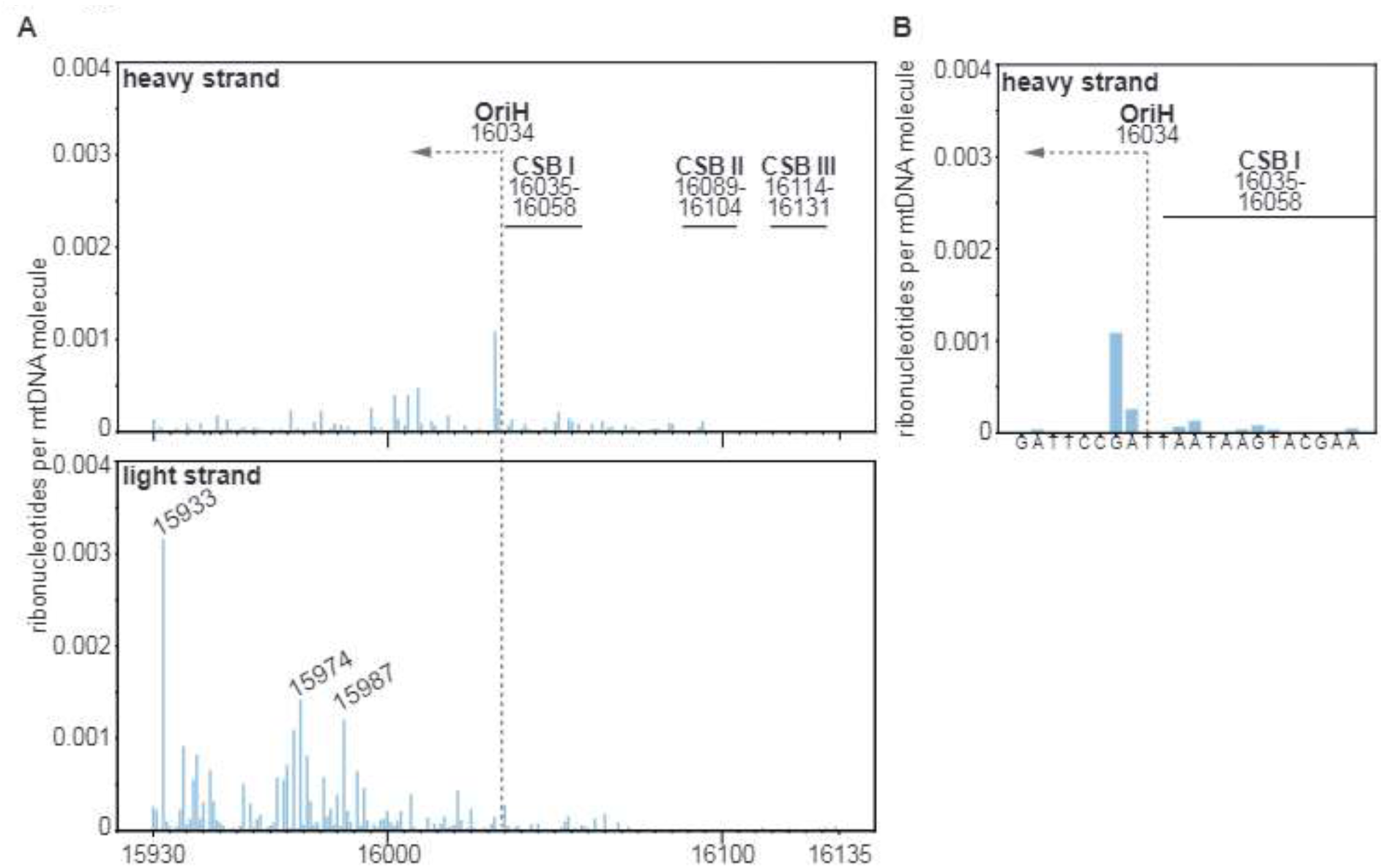
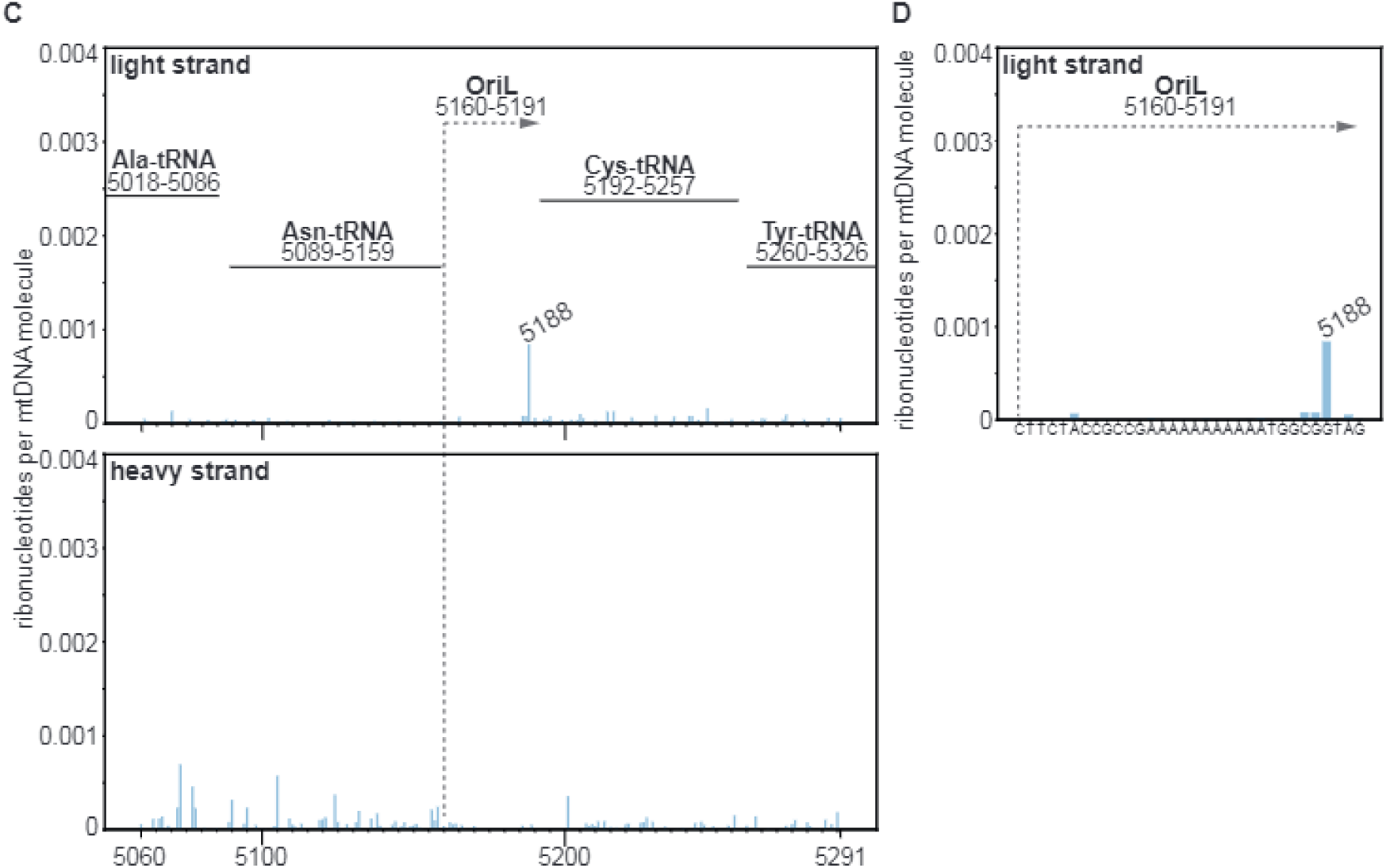
Ribonucleotide hotspots near OriH and OriL. **A** Ribonucleotides per heavy or light strand near OriH. **B** Close view of the ribonucleotide hotspot at 16,032 near OriH. **C** Ribonucleotides per light or heavy strand near OriL. **D** Close view of the ribonucleotide hotspot at 5,188 near OriL. (All tissues, n = 51)

## DISCUSSION

Since incorporated ribonucleotides are estimated to be the most common non-canonical nucleotides in eukaryotic DNA^9^, that they are much more unstable than deoxyribonucleotides^13^ and have the potential to alter the DNA structure^41,42^, these ribonucleotides can have significant consequences for genome stability. Their distribution in the mammalian genome and in relation to common genomic features are the first steps to precisely discern what determines their appearance, permanent incorporation, recognition or protection from removal mechanisms and the resulting beneficial or detrimental effects of their presence. Here we have for the first time determined the number and identity of incorporated ribonucleotides in wild-type mouse nDNA and mtDNA across nine tissues: blood, bone marrow, brain, heart, kidney, liver, lung, muscle and spleen. The number of incorporated ribonucleotides and their base identity varied between the nDNA of the investigated tissues and to a greater extend also in mtDNA. Our estimate of about 5.2 million stably incorporated ribonucleotides per murine nuclear genome suggests that ribonucleotides are the most common non-canonical nucleotides in DNA. This number is higher than anticipated from RNase H2 deficient murine embryonic fibroblasts (MEFs) where an incorporation of 1 ribonucleotide per 7.6 kb nucleotides equivalent to about 1.3 million ribonucleotides per genome were estimated to be incorporated and a near complete repair was expected^43^. Several factors may contribute to different estimates in the number of incorporated ribonucleotides between MEFs and the investigated mouse tissues: differences between tissues (Fig. 1 A and B) and that our tissues were obtained from 7.5 or 30 week-old mice rather than embryos allowing an accumulation over time are conceivable contributing factors. Furthermore, the incorporated ribonucleotides may be the result of the cell cycle-dependent regulation of RNase H2, which seems to limit its activity to the G2 phase^44^. Ribonucleotide removal efficiency might be further restricted by impeded recognition of incorporated ribonucleotides through masking by other DNA-interacting or -binding proteins, as suggested by our findings. While the differences between nDNA and mtDNA are most likely explained by the lack of efficient ribonucleotide removal in mitochondria^25,29^ and differences in sugar selectivity between the replicative DNA polymerases and mtDNA polymerase γ^8^, the tissue-specific differences can probably be attributed to factors determined by the specific characteristics of the cell types of each given tissue. This includes for example when and how cells undergo cell cycles^45^ which in turn regulate dNTP availability through cell cycle-controlled expression of ribonucleotide reductase^46^, thymidine kinase^47^ and the dNTP triphosphohydrolase, sterile alpha motif domain and histidine-aspartate domain-containing protein 1 (SAMHD1)^48^ and ribonucleotide removal by regulating the expression of RNase H2^44,49^. As a cell organelle separated from the cytosol, additional regulatory mechanisms affect the available NTP pools in mitochondria. Mitochondrial dNTPs underly a complex cross-talk between the cytoplasm and the mitochondria and can either be produced by salvage pathway enzymes or are imported from the cytoplasm via mitochondrial nucleotide transporters^50^. Mitochondrial NTP pools directly affect the number and identity of incorporated ribonucleotides in mtDNA^25^ and loss of SAMHD1, which can hydrolyze all four dNTPs in the presence of deoxyguanosine triphosphate^51^ leads to a low frequency of ribonucleotides in mouse mtDNA^26^.

Dysregulation or defects of factors involved in NTP pool maintenance and ribonucleotide removal and resulting imbalance in NTP pools and ribonucleotide accumulation in DNA are of interest because of their association with human disease and models for human disease: among others, mutations in SAMHD1, and the RNase H2 subunits, RNASEH2A, RNASEH2B, RNASEH2C can cause Aicardi-Goutières syndrome, an inflammatory encephalopathy^52^ or systemic lupus erythematosus, which shares similarities with the Aicardi-Goutières syndrome^53^. RNase H2 deficiency in the epidermis of mice was found to increase spontaneous DNA damage and skin cancer^15^. Epithelial RNase H2 was also shown to be important in the prevention of intestinal tumorigenesis in mice^54^. Acetylated SAMHD1, which has increased dNTPase activity, is implicated in promoting cancer cell proliferation in both tumor cells from hepatocarcinoma patients and HeLa cells^55^. CRISPR screening identified TOP1-incised embedded ribonucleotides as poly(ADP-ribose) polymerase (PARP) trapping lesions that impede DNA replication and affect PARP inhibitor cytotoxicity^56^. This might be exploited to sensitize cells in the treatment of prostate cancer and chronic lymphocytic leukemia which are frequently deficient in RNASEH2B^57^. The accumulation of incorporated ribonucleotides in the absence of RER can furthermore lead to unrepaired nicks that may contribute to the pathology of ataxia with oculomotor apraxia 1^58^. Some pathologies are more specific to the mitochondrial NTP pools and the mtDNA integrity in particular. Several forms of mtDNA depletion syndromes can be caused by the proteins involved in maintaining mitochondrial dNTP pools, their disruption leading to limited dNTPs available to mtDNA replication^59^. Mutations in RNASEH1, which is responsible for the primer removal in mtDNA^37^, can cause adult-onset mitochondrial encephalomyopathy^60^.

Our meta-analyses of genomic features imply that the non-random distribution of ribonucleotides in the murine nDNA is probably the product of a variety of processes affecting incorporation and removal dependent of the genomic feature. Because cleavage of a RNA-DNA junction for ribonucleotide removal requires access of the enzymes’ active site^61^ it seems reasonable to suspect that DNA binding proteins or other DNA-interacting proteins, as well as structural distortion in the case of G4s (Fig. 3 B) may mask incorporated ribonucleotides from recognition by RNase H2 or TOP1, but mechanistic experiments are needed to test this hypothesis.

While an accumulation of incorporated ribonucleotides seems to generally have negative effects as described above, also more studies emerge that suggest beneficial roles of transiently incorporated ribonucleotides involved in DNA repair intermediates, illustrating the balancing act between advantageous and detrimental ribonucleotide incorporation. Newly embedded ribonucleotides were found to serve as a strand discrimination signal in eukaryotic mismatch repair and mismatch repair efficiency is decreased in the absence of RNase H2^11^. In the repair of double-strand breaks by mammalian non-homologous end-joining, ribonucleotides incorporated by DNA polymerase μ or terminal deoxynucleotidyl transferase were shown to facilitate first-strand ligation in as many as 65% of non-homologous end-joining products^62,63^. Moreover, recent findings suggest a RNA-DNA hybrid produced by RNA polymerase III to be an important intermediate for the protection of the 3’-single-stranded DNA overhang in homologous recombination, thereby explaining the need for RNase H1 and 2 for efficient double-strand break repair by homologous recombination^64^. RNase H2’s cellular significance is further supported by the marked transcriptional changes in the cell, including the increase in expression of genes involved in nucleic acid transactions, helicases and nucleases in the absence of RNH201 in yeast^65^.

Ribonucleotides embedded in the mtDNA seem to be tolerated better than in nDNA, as we observed a higher frequency of ribonucleotides in mtDNA of wild-type mice (Fig. 1 B) and given the fact that mtDNA stability does not seem to be negatively impacted by the age-related accumulation of ribonucleotides in mtDNA due to the lack of efficient ribonucleotide removal^26^. It is therefore of interest to understand if their presence contributes to mtDNA instability in other ways. In the mouse model for a mitochondrial DNA depletion syndrome caused by MPV17 deficiency, mtDNA deletions and an increased number of incorporated rG were observed in the brains of MPV17 knockout mice. It was therefore hypothesized that those ribonucleotides may be involved in causing these deletions^16,40^. Interestingly, mutational hotspots identified by ultra-deep sequencing in mouse mtDNA only coincided with the ribonucleotide peak we observed near OriL at 8,155 (Fig. 5 D), while none of the ribonucleotide peaks listed in Figure 4 matched the most frequently mutated areas of the mitochondrial genome^66^, which could imply that embedded ribonucleotides in mtDNA are not the predominant cause for arising mutations.

Despite their relevance to human health and the surge of interest in incorporated ribonucleotides in nDNA and mtDNA, their implications in maintaining or hurting genome integrity, their precise physiological roles remain yet to be fully understood.

## MATERIALS AND METHODS

### Mice

All procedures were performed in accordance with the institutional guidelines of the University of Gothenburg and experiments were approved by the Institutional Review Board of the Swedish Board of Agriculture (Ethical approval number 62/14 and 1469/2019). Samples from three 7.5-week-old and three 30-week-old male mice (C57BL6) were used for our experiments. Mice were euthanized by placing them into an induction chamber (Abbott Scandinavia AB) for 10 min with 4% isoflurane (Forene isoflurane, Abbott Scandinavia AB) followed by cervical dislocation. Whole blood was immediately harvested and the heart surgically excised thereafter. 300-600 µL of whole blood were thoroughly mixed with 30 µl 0.1 M EDTA to prevent coagulation, 200 µL aliquots were transferred to 1.5 mL tubes and frozen at -80 °C. Tibia and femur bones (for extraction of bone marrow), brain, heart, kidney, liver, lung, thigh muscle and spleen were removed and small sections were flash frozen with liquid nitrogen, then stored at -80 °C until further processing.

### DNA extraction

DNA was extracted using the MasterPure Complete DNA and RNA purification kit (MC85201, lucigen) preceded by adjusted approaches for homogenization and lysis of tissues. DNA from whole blood was extracted using the kit following manufacturer’s instruction for the extraction from whole blood with RBC lysis. DNA from bone marrow originated from tibia and fibula. Bones were freed from any remaining tissue and broken at the knee joints. Placing this opening downward in a perforated 0.5 mL tube set in a 1.5 mL tube allowed to centrifuge out the bone marrow at 10,000 g for 20 s. 100 µL of RCB lysis buffer were added and mixed and then incubated at RT for 10 min while vortexing every 5 min. Samples were centrifuged at 10,000 g for 25 s and supernatant discarded. 300 µL of tissue and cell lysis buffer including proteinase K were added and incubated at 65 °C for 15 min while shaking at 1,500 rpm. Further extraction steps were performed following the kit’s instructions. 2-70 mg of brain, heart, kidney, liver, lung, muscle or spleen tissue were added to a 1.5 mL tube containing 5-15 stainless steel homogenizing beads (0.9-2 mm, SSB14B, next advanced Inc.) and 100 µL tissue and cell lysis buffer including proteinase K. Samples were vortexed for about 20-30 min until tissue pieces were no longer visible. Beads were pelleted using a magnetic rack (DynaMag-2, Invitrogen) and the supernatant was transferred to a new tube containing 200 µL of tissue and cell lysis buffer including proteinase K. Samples were incubated at 65 °C for 15 min while shaking briefly every 5 min. After cooling the samples on ice for 5 min, further extraction steps were performed as instructed. Final DNA concentrations were determined using the Qubit dsDNA BR Assay Kit (molecular probes life technologies).

### Mapping and quantitation of 5’-ends and ribonucleotides

500 ng to 1 μg gDNA were cleaved by SacI-HF (New England Biolabs) for 15 min at 37 °C. The reaction was stopped by bead washing using 1.8 volumes CleanPCR paramagnetic beads (CleanNA) following manufacturer’s instructions. The gDNA was eluted in EB and used as input for the HydEn-seq method as described previously^27,67^. In brief, alkaline hydrolysis was performed on the cleaved gDNA using 0.3 M KOH (or 0.3 M KCl for background controls) for 2 h at 55 °C. Resulting DNA fragments were precipitated with ethanol, denatured and phosphorylated with 10 U of 3’-phosphatase-minus T4 polynucleotides kinase (New England Biolabs) at 37 °C for 30 min. The reaction was terminated by heat inactivation at 65 °C for 20 min. After DNA purification using the CleanPCR beads, the DNA was denatured and ligated to oligo ARC140 overnight at room temperature by 10 U T4 RNA ligase (New England Biolabs) in 25% PEG8000 and 1 mM CoCl_3_(NH_3_). DNA was purified using CleanPCR beads and denatured before the ARC76-66 adaptor was annealed at room temperature for 5 min. 4 U of T7 DNA polymerase (New England Biolabs) were used to perform second-strand synthesis and DNA was purified using CleanPCR beads. Librarie were amplified using individual index primers ARC79 to ARC107 by the KAPA Hifi Hotstart ReadyMix (KAPA Biosystems). CleanPCR beads were used to purify the libraries before pooling them for sequencing. An Illumina NextSeq500 instrument was used for 75-bp paired-end sequencing, to locate the 5’-ends resulting from the alkaline hydrolysis. Each library corresponded to one distinct tissue sample and replicates for each tissue came from different mice (n = 6 for blood, brain, heart, kidney, liver and muscle; n=5 for bone marrow, lung and spleen). All tissue samples were summarized in the analyses for Figures 3 to 5 (n = 51).

### Sequence trimming, filtering and alignment

Cutadapt 1.12^68^ was used for quality and adaptor sequence trimming. Pairs with one or both reads shorter than 15 nt were discarded. Mate 1 of the remaining pairs was aligned to the list of index primers used to prepare the libraries using Bowtie 1.2 and all matching pairs were removed. Remaining pairs were aligned to the mm10/GRCm38 *Mus musculus* reference genome (https://www.ncbi.nlm.nih.gov/grc) using bowtie (-v2 –X2000-best). Then, single-end alignments were performed for mate 1 of all unaligned pairs (-m1, -v2). The count of 5’-ends of all unique paired-end and single-end alignments was determined and shifted one base upstream to represent the location of the original embedded ribonucleotide. For the mtDNA, reads were uniquely aligned only to the mitochondrial chromosome.

### Quantitation of incorporated ribonucleotides

For comparison between libraries, visual representations, and meta-analyses all end counts were normalized to the mean reads per SacI cleavage site detected in each sample. Normalized 5’-end counts from KCl-treated control samples were subtracted from the corresponding KOH-treated samples to remove counts from free 5’-ends that were not caused by alkaline hydrolysis.

### Analysis of ribonucleotides near genomic features

A list of confirmed mouse TSS was retrieved from the RefTSS data set (http://reftss.clst.riken.jp/reftss/)^36^. Positions of intrastrand G4s were predicted by using a custom script searching the mm10 reference genome for the motif (G_3+_N_1-25_)_3_G_3+_. Other genomic positions for ROI were acquired from the UCSC table browser (https://genome.ucsc.edu/cgi-bin/hgTables)^35^: enhancer^30,69^, tRNAs^31^, promoters^32^, CpG islands^33^ and microsatellites^34^. For the calculation of ribonucleotide enrichment or depletion, we generated a data set of 21,000 random genomic positions by picking 1,000 random positions from each chromosome. Data processing and visualization was performed using custom scripts: All 5’-end counts were normalized to the mean reads per SacI cleavage site and normalized end counts from KCl-treated samples were subtracted from the corresponding SacI-normalized end counts of KOH-treated samples to obtain only reads at incorporated ribonucleotides. All reads for ribonucleotides were collected near each ROI start position including a 10kb, 4 kb, 1 kb or 500 b window up- and downstream of the ROI and binned in 500 b-, 200 b-, 50 b- or 25 b-bins, respectively. The mean number of ribonucleotides per bin was calculated. The same procedure was performed with the random data set, which was then used to calculate ribonucleotide enrichment.

Similarly, mtDNA visualization (Fig. 4 and 5) was performed by normalizing all reads to mean reads per SacI cleavage site and subtracting values from KCl-treated libraries from the corresponding KOH-treated libraries to calculate the mean number of ribonucleotides on each position of a mtDNA molecule of all samples. Whole mtDNA overviews of mean ribonucleotides per heavy or light strand of one mtDNA molecule position were displayed using Circos^70^. Visualizations of 5’-ends on mtDNA heavy and light strand resulting from KCl or KOH treatment normalized to mean reads in SacI sites are provided in Supplementary Figure 1.

---

**Supplementary Figure 1.**
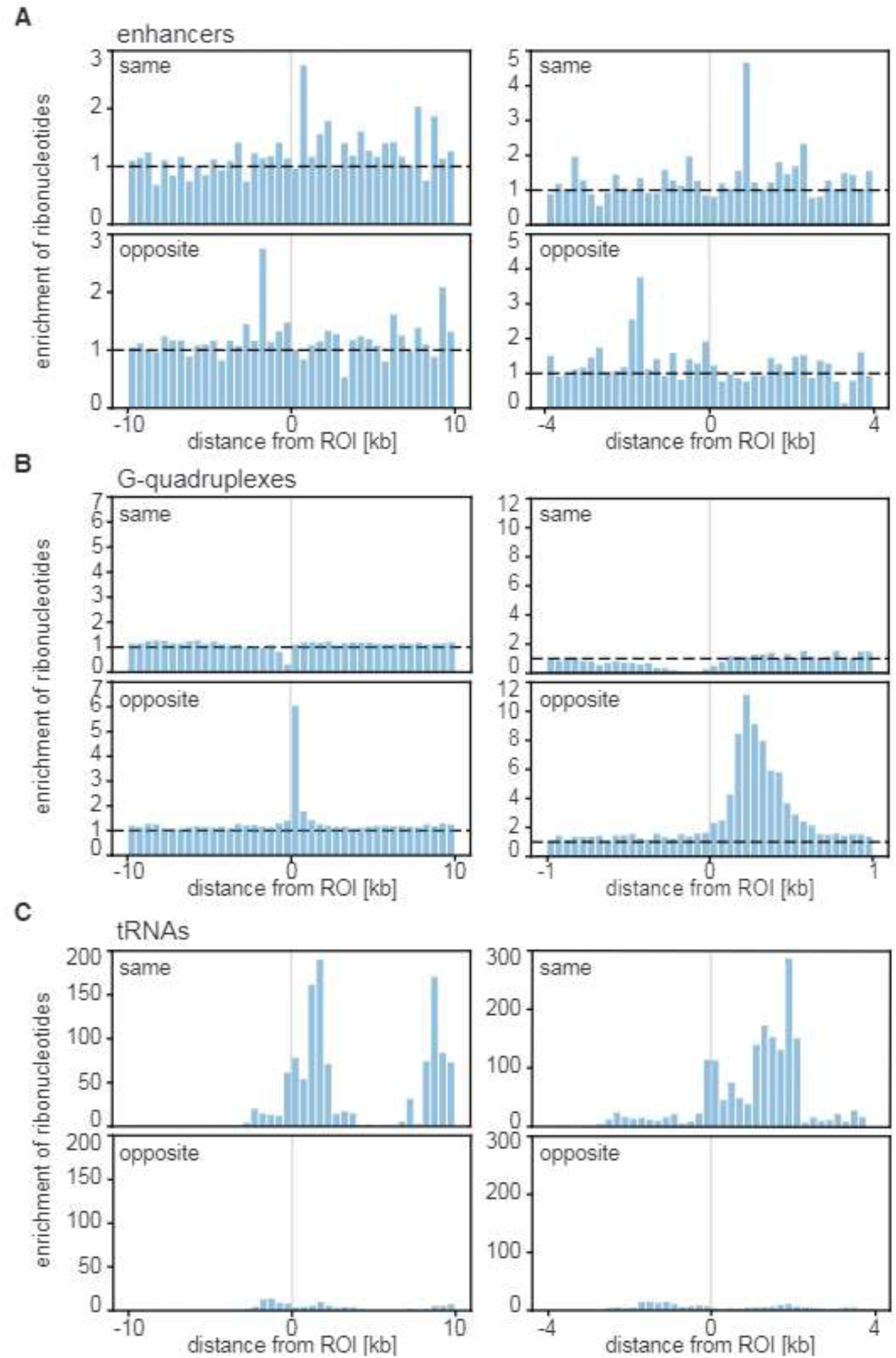
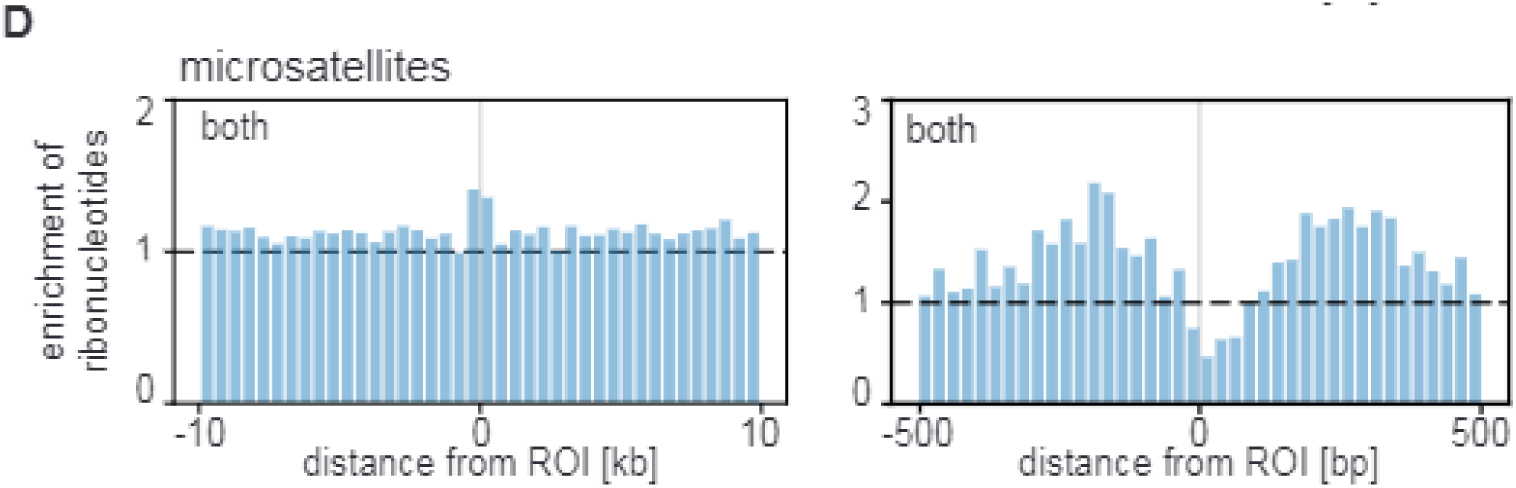
Enrichment of ribonucleotides at genomic features. Enrichment of ribonucleotides near genomic region of interest (ROI) on same, opposite or both strands. **A** Enrichment of ribonucleotides near 863 enhancers on the same and opposite strands. Mean ribonucleotide enrichment per 500 b-bin in a 10 kb-window (left panels) and per 200 b-bin in a 4 kb-window (right panels) up- and downstream of the ROI are shown. **B** Enrichment of ribonucleotides near 1,048,573 intrastrand G-quadruplexes on the same and opposite strands. Mean ribonucleotide enrichment per 500 b-bin in a 10 kb-window (left panels) and per 50 b-bin in a 1 kb-window (right panels) up- and downstream of the ROI are shown. **C** Enrichment of ribonucleotides near 435 tRNAs on the same and opposite strands. Mean ribonucleotide enrichment per 500 b-bin in a 10 kb-window (left panels) and per 200 b-bin in a 4 kb-window (right panels) up- and downstream of the ROI are shown. **D** Enrichment of ribonucleotides near 197,237 microsatellites on both strands. Mean ribonucleotide enrichment per 500 b-bin in a 10 kb-window (left panels) and per 25 b-bin in a 500 bp-window (right panels) up- and downstream of the ROI are shown. (All tissues, n = 51)

## ACKNOWLEDGMENTS

We thank the Genomics Core Facility at the Sahlgrenska Academy, University of Gothenburg for sequencing. This study was supported by the Swedish Research Council (www.vr.se) [2018-05121 to A.R.C.].

## CONFLICT OF INTEREST

The authors declare that they have no conflicts of interest with the contents of this article. J.K. is an employee of Astra Zeneca at the time of publishing, but the work presented in this manuscript is unrelated and was performed prior to this affiliation.

## DATA AVAILABILITY

The datasets generated and analysed during the current study are available in the GEO repository, accession number GSE183589 (https://www.ncbi.nlm.nih.gov/geo/query/acc.cgi?acc=GSE183589).

## CODE AVAILABILITY

Code is available upon request.

## AUTHOR CONTRIBUTIONS

A.R.C. designed experiments. L.M.A. provided laboratory mice. S.B. and C.A. sacrificed the mice and harvested all tissues. J.K. performed experiments. K.K. and A.R.C. analyzed data. K.K. wrote the manuscript. K.K. and A.R.C. revised the manuscript with input from all authors.

## Notes

### Competing Interest Statement

The authors have declared no competing interest.

